# A chromosome-level, fully phased genome assembly of the oat crown rust fungus *Puccinia coronata* f. sp. *avenae*: a resource to enable comparative genomics in the cereal rusts

**DOI:** 10.1101/2022.01.26.477636

**Authors:** Eva C. Henningsen, Tim Hewitt, Sheshanka Dugyala, Eric S. Nazareno, Erin Gilbert, Feng Li, Shahryar F. Kianian, Brian J. Steffenson, Peter N. Dodds, Jana Sperschneider, Melania Figueroa

## Abstract

Advances in sequencing technologies as well as development of algorithms and workflows have made it possible to generate fully phased genome references for organisms with non-haploid genomes such as dikaryotic rust fungi. To enable discovery of pathogen effectors and further our understanding of virulence evolution, we generated a chromosome-scale assembly for each of the two nuclear genomes of the oat crown rust pathogen, *Puccinia coronata* f. sp. *avenae* (*Pca*). This resource complements two previous released partially phased genome references of *Pca*, which display virulence traits absent in the isolate of historic race 203 (isolate *Pca*203) which was selected for this genome project. A fully phased, chromosome-level reference for *Pca*203 was generated using PacBio reads and Hi-C data and a recently developed pipeline named NuclearPhaser for phase assignment of contigs and phase switch correction. With 18 chromosomes in each haplotype and a total size of 208.10 Mbp, *Pca*203 has the same number of chromosomes as other cereal rust fungi such as *Puccinia graminis* f. sp. *tritici* and *Puccinia triticina*, the causal agents of wheat stem rust and wheat leaf rust, respectively. The *Pca*203 reference marks the third fully-phased chromosome-level assembly of a cereal rust to date. Here, we demonstrate that the chromosomes of these three *Puccinia* species are syntenous and that chromosomal size variations are primarily due to differences in repeat element content.

## Introduction

The rust fungus *Puccinia coronata* f. sp. *avenae* (*Pca*) is the most damaging foliar pathogen of oat (*Avena sativa* L.) (Nazareno *et al.* 2018). *Pca* is commonly found in areas of the world where oats are grown and can destroy up to 50% of the crop during severe epidemics (USDA-ARS CDL 2014; Nazareno *et al.* 2018). Plants have race-specific resistance (*R*) genes, which generally encode immunoreceptors that recognize pathogen effectors as a strategy to stop infection and disease development (Dodds and Rathjen 2010). The molecular basis of virulence in *Pca* is not well characterized and this is partly due to the lack resources to direct such studies. High quality genome references of the pathogen are instrumental to investigate the underlying mechanisms of virulence, enable effector discovery, and even accelerate the identification of *R* genes (Figueroa *et al*. 2016, 2020; Upadhyaya *et al*. 2021).

The dikaryotic nature of rust fungi means their genetic information is present in two haplotypes that are physically separated in two nuclei for most of their life cycle. This feature in combination with highly repetitive genomes poses some technical challenges for genome assembly. Until recently, genome references built for rust fungi did not fully capture sequence information from both nuclei and most assemblies were haploid representations of the genome with abundant haplotype sequence (phase) swaps (Cantu *et al*. 2011; Cuomo *et al*. 2017). New technologies such as long-read and Hi-C sequencing (Li *et al*. 2019) as well as the recent development of the NuclearPhaser pipeline (Duan *et al*. 2021) have improved our ability to assemble chromosome references and fully resolve the two haplotypes in rust fungi. To date the only existing chromosome-level nuclear phased assemblies belong to *Puccinia graminis* f. sp. *tritici* and *Puccinia triticina*, which cause stem and leaf rust on wheat, respectively. There are two *Pca* genome assemblies (Miller *et al*. 2018), which were the first attempt towards haplotype resolution of any rust fungi and consist of a primary pseudo-haplotype assembly, with a partially resolved secondary haplotype that is only 50-60% complete.

Here, we selected another *Pca* isolate, *Pca*203, for assembly of a complete haplotype-phased genome reference using state-of-art *de novo* genome assembly approaches for dikaryotic rust fungi. *Pca*203 represents an isolate of the historic *Pca* race 203, which caused severe epidemics in the early 1940s in the US (Stoa and Swallers, 1950) leading to the deployment of the *Pc2* resistance gene present in Victoria oats. The subsequent widespread cultivation of Victoria oats led to devastating epidemics of *Cochliobolus victoriae*, likely as a result of *Pc2* acting as a susceptibility factor for the Victorin toxin produced by this necrotrophic pathogen (Murphy and Meehan 1946; Wolpert et al., 2002). Thus, a gold-standard assembly of *Pca*203 could bring insights into one of the best-known classical problems of plant pathology. Furthermore, the contrasting virulence profile of *Pca*203 and other previously sequenced *Pca* isolates will help unravel the molecular basis of *Pca* virulence and aid in future comparative genomic studies within *Pca* and other rust species.

## Materials and Methods

### Plant and fungal materials and plant inoculations

An isolate of *Pca* race 203, known to be avirulent to the oat cultivar Victoria (Chang and Sadanaga 1964), was retrieved from storage at the USDA-ARS Cereal Disease Laboratory (CDL), Saint Paul, MN, U.S.A (Omidvar et al., 2018). From this culture, a single pustule was isolated and increased to ensure sample purity. The virulence phenotype of *Pca*203 was determined according to a current standard nomenclature system using a set of oat differentials (Chong et al. 2000; Nazareno et al. 2018). Urediniospore stocks were kept at −80° C. As previously described by Miller et al. (2018), virulence phenotypes of *Pca* on the oat differentials were converted to a 0-9 numerical scale for heat map generation using R packages ComplexHeatmap and circlize (Gu et al. 2016, 2014). Seed from the oat differential set was also obtained from the USDA-ARS CDL. Urediniospores increases were completed on the oat cv. Marvelous as a susceptible host. Oat inoculations with *Pca* were carried out as described by Omidvar et al. (2018).

### DNA and RNA isolation and sequencing

High molecular weight DNA was extracted from 700 mg of urediniospores, as previously described by Li et al. (2019) and sent to the University of Minnesota Genomics Center (UMGC, St. Paul, MN, U.S.A.) for library construction using the PacBio SMRTbell 1.0 kit and sequencing using five PacBio Sequel System SMRT cells with v3 chemistry. DNA was also extracted from 20 mg of isolate *Pca*203 spores using the Omniprep™ DNA isolation kit from G-Biosciences for library preparation with the Illumina TruSeq Nano DNA protocol and Illumina NovaSeq sequencing in the S2 flow cell at UMGC to produce 150 bp paired-end reads. For DNA-crosslinking and Hi-C sequencing, 100 mg of spores were suspended in 15 mL 1% formaldehyde and incubated at room temperature (RT) for 20 minutes with periodic vortexing. Glycine was added to 1g/100mL, and the suspension was incubated again at RT for 20 minutes with periodic vortexing. The suspension was centrifuged at 1000*g* for 1 minute and the supernatant was removed; the spores were then transferred to a liquid nitrogen-cooled mortar and ground before being stored at −80° C or on dry ice. Crosslinked spores were sent to Phase Genomics (Seattle, WA, U.S.A) for Hi-C library preparation with the Proximo Fungal 4.0 protocol and libraries were sequenced to 100 million 150 bp paired-end reads at Genewiz (South Plainfield, NJ, U.S.A.). RNA was extracted from infected oat cv. Marvelous at two- and five-days post-inoculation (dpi) using the Qiagen RNeasy Plant Mini Kit following the manufacturer’s instructions and sent to UMGC for library preparation with the Illumina TruSeq Stranded mRNA protocol and sequencing using Illumina NextSeq in mid-output mode, producing 75 bp paired-end reads.

### Genome assembly and polishing

Rust isolate purity was assessed by examining SNP allele balance as previously described (Miller et al. 2018). For this, Illumina short reads were trimmed with trimmomatic version 0.33 (Bolger et al. 2014) and aligned with bwa version 0.7.17 (Li and Durbin 2009) to the existing 12SD80 primary genome assembly (Miller et al. 2018). The alignments were processed using samtools version 1.9 (Li et al. 2009), and variants were called with Freebayes version 1.1.0 (Garrison and Marth 2012). The allele balance plot was generated using a custom R script (https://github.com/henni164/Pca203_assembly/figure_s1/203_frequencies.R).

The genome was assembled using Canu version 2.1 with setting genomeSize=200m (Koren et al. 2017). For polishing with the PacBio read data, half of the subreads were mapped back to the assembly using PacBio software pbmm2 version 1.4.0 (https://github.com/PacificBiosciences/pbmm2/releases/tag/v1.4.0) and a new consensus generated with PacBio software GenomicConsensus version 2.3.3 (https://github.com/PacificBiosciences/GenomicConsensus/releases/tag/2.3.3). This process was repeated using the new consensus as the reference. Next, the updated consensus was polished twice in Pilon version 1.22 (--fix indels) using Illumina short reads trimmed with trimmomatic version 0.33 (Bolger et al. 2014; Walker et al. 2014). Assembly statistics were calculated using Quast version 5.1.0 (Gurevich et al. 2013).

### Identification of mitochondrial contigs and removal of contaminants

All contigs were first screened by BLAST against the mitochondrial genome database from NCBI with ncbiblast+ version 2.8.1, and mitochondrial contigs were removed from the main assembly (Camacho et al. 2009). The remaining contigs were screened against the NCBI nucleotide library and eight other likely contaminant contigs were removed. Finally, eight contigs with PacBio reads coverage <2x were removed. Telomeres and collapsed regions were identified with custom scripts (https://github.com/JanaSperschneider/GenomeAssemblyTools, https://github.com/JanaSperschneider/FindTelomeres). One contig consisting entirely of telomeric repeat reads was also removed.

### Haplotype-phasing and annotation of genome assembly

The NuclearPhaser pipeline was used to correct phase swaps and assign haplotypes as described by Duan et al. (2021), except that two rounds of phase swap correction instead of one round were conducted. Likely phase swap locations were located by plotting the proportion of Hi-C *trans-* contacts to the partially phased haplotypes, as described in https://github.com/JanaSperschneider/NuclearPhaser. Phase swap breakpoints were identified by examining short read alignments in Integrative Genomics Viewer (IGV) and choosing positions with either high coverage of multimapping reads (representing highly similar regions which were phased, but assembled into the wrong haplotype) or high coverage of unique mapping reads with high SNP density (representing collapsed regions). Collapsed regions were included in both haplotypes.

For scaffolding of the phased haplotypes, the Hi-C reads were mapped to each haplotype using BWA-MEM version 0.7.17 (Li and Durbin, 2009) and alignments were then processed with the Arima Genomics pipeline (https://github.com/ArimaGenomics/mapping_pipeline/blob/master/01_mapping_arima.sh). Scaffolding was performed using SALSA version 2.2 (Ghurye *et al*., 2017, 2019).

*De novo* repeats were predicted with RepeatModeler 2.0.0 and the option -LTRStruct (Flynn *et al*., 2020). The predicted repeats were merged with the RepeatMasker repeat library and RepeatMasker 4.1.0 was run with this combined repeat database (http://www.repeatmasker.org). The resulting repeat-masked genome was used for gene annotation. RNAseq reads were cleaned with fastp 0.19.6 using default parameters (Chen *et al*., 2018). RNAseq reads were aligned to the genome with HISAT2 (version 2.1.0 -max-intronlen 3000 – dta) (Kim *et al*., 2019). Genome-guided Trinity (version 2.8.4 –jaccard_clip – genome_guided_bam – genome_guided_max_intron 3000) was used to assemble transcripts (Grabherr *et al*., 2011). Annotation was performed with funannotate version 1.7.4 (Palmer and Stajich 2020). First, funannotate train was run with the Trinity transcripts. Second, funannotate predict was run on the repeat-masked genome with options –ploidy 2 –optimize_augustus. Third, funannotate update was run (--jaccard_clip). BUSCO scores were obtained by running BUSCO version 3.1.0 (Waterhouse *et al*. 2018). Secreted proteins were predicted with SignalP 4.1 (-t euk -u 0.34 -U 0.34) (Petersen *et al*. 2011) and TMHMM version 2.0 (Krogh *et al*. 2001). A fungal protein was called secreted if it was predicted to have a signal peptide and has no transmembrane domains. Effector proteins were predicted with EffectorP version 3.0 (Sperschneider and Dodds 2021).

Hi-C contact maps were produced with HiC-Pro 2.11.1 (MAPQ=10) (Servant *et al*., 2015) and Hicexplorer 3.6 (Ramírez *et al*., 2018; Winter *et al*., 2018; Wolff *et al*., 2018, 2020). The normalized Hi-C contact maps were used to plot the distribution of Hi-C links within and between the two haplotypes.

### Comparative genomics and phylogenetic analysis

The *Pca*203 genome was aligned to the *Pgt*21-0 and *Pt*76 genomes using D-Genies with the minimap2 alignment setting (Cabanettes *et al*. 2018; Li *et al*. 2018, Duan *et al*. 2021). Phylogenetic analyses were performed on a set of 63 isolates: 30 isolates from 1990, 30 isolates from 2015 as published by Miller et al. (2020), and the three genome reference representatives (12SD80, 12NC29 and *Pca*203). Trimmed Illumina reads were aligned to the 12SD80 reference primary contigs using bwa version 0.7.17 (Li and Durbin 2009). The resulting BAM files were processed with Samtools version 1.9 (Li et al. 2009) including removal of duplicate reads and filtering for mapping quality of 30. Variant calling against the 12SD80 reference primary contigs was performed using Freebayes version 1.3.2. The resulting VCF file was filtered with vcffilter in vcflib version 1.0.1 (https://github.com/vcflib/vcflib) using the parameters “QUAL > 20 & QUAL / AO > 10 & SAF > 0 & SAR > 0 & RPR > 1 & RPL > 1 & AC > 0”. Additional filtering for 90% genotyping frequency (<10% missing data), 5% minor allele frequency and for bi-allelic SNPs was performed using vcftools version 0.1.16 (Danecek *et al*. 2011) to give a final VCF file representing 974,924 variant sites. To analyze reticulation in *Pca*, an unrooted phylogenetic network was created using SplitsTree version 4.16.2 (Huson and Bryant 2006). The network tree was exported in scalable vector graphics (SVG) format and labels modified using Inkscape version 1.1.1 (https://inkscape.org/).

### Data Availability

Sequencing reads, the assembly and annotation are available at the CSIRO Data Portal https://data.csiro.au/collection/csiro:53477. Scripts for identification of contaminants, collapsed regions, and telomeres are available at https://github.com/JanaSperschneider/GenomeAssemblyTools and https://github.com/JanaSperschneider/FindTelomeres. NuclearPhaser is available at https://github.com/JanaSperschneider/NuclearPhaser. Scripts and data used to construct figures are available at https://github.com/henni164/Pca203_assembly. Sequencing data from the SRA numbers listed in **Table S1**.

## Results and Discussion

### Virulence profile and pathotyping of *Pca* isolate 203

To create a chromosome-level nuclear phased assembly, we selected isolate *Pca*203, which represents the race of the oat crown rust pathogen, named 203. The *Pca* race 203 was very common in North America in the 1940s and was still present in the 1970’s (Fleischmann, 1967; Fleischmann and Baker 1971, Stoa and Swallers, 1950). The isolate *Pca*203 was previously revived from a long-term storage collection at the USDA-ARS CDL (Omidvar et al., 2018). Here, we reanalyzed the virulence profile and adopted the current pathotype nomenclature to characterize of the isolate *Pca*203 stocks of urediniospores. A full report of infection scores of *Pca*203 with the current expanded oat differential set that is currently used in North America (Nazareno *et al*. 2018) was generated (**Figure 1A**) and compared to previously sequenced isolates (Miller et al, 2018; 2020). *Pca*203 was assigned to pathotype BBBGBCGLLB. Compared to the *Pca* isolates (12NC29 and 12SD80) for which genome references have been previously constructed (Miller *et al*. 2018), *Pca*203 has a unique virulence profile as most *R* genes confer resistance against it (**Figure 1**). This suggests that *Pca*203 contains Avr effectors that are absent in 12NC29 and 12SD80 and the *Pca*203 genome can therefore assist in effector identification and studies of virulence evolution for resistance genes represented in the oat differential set. A full comparison to the virulence profiles of isolates derived from 2015 and 1990 (Miller *et al*. 2020) is shown in **Figure S1** to demonstrate the abundance of avirulence traits in *Pca*203.

**Figure 1.**
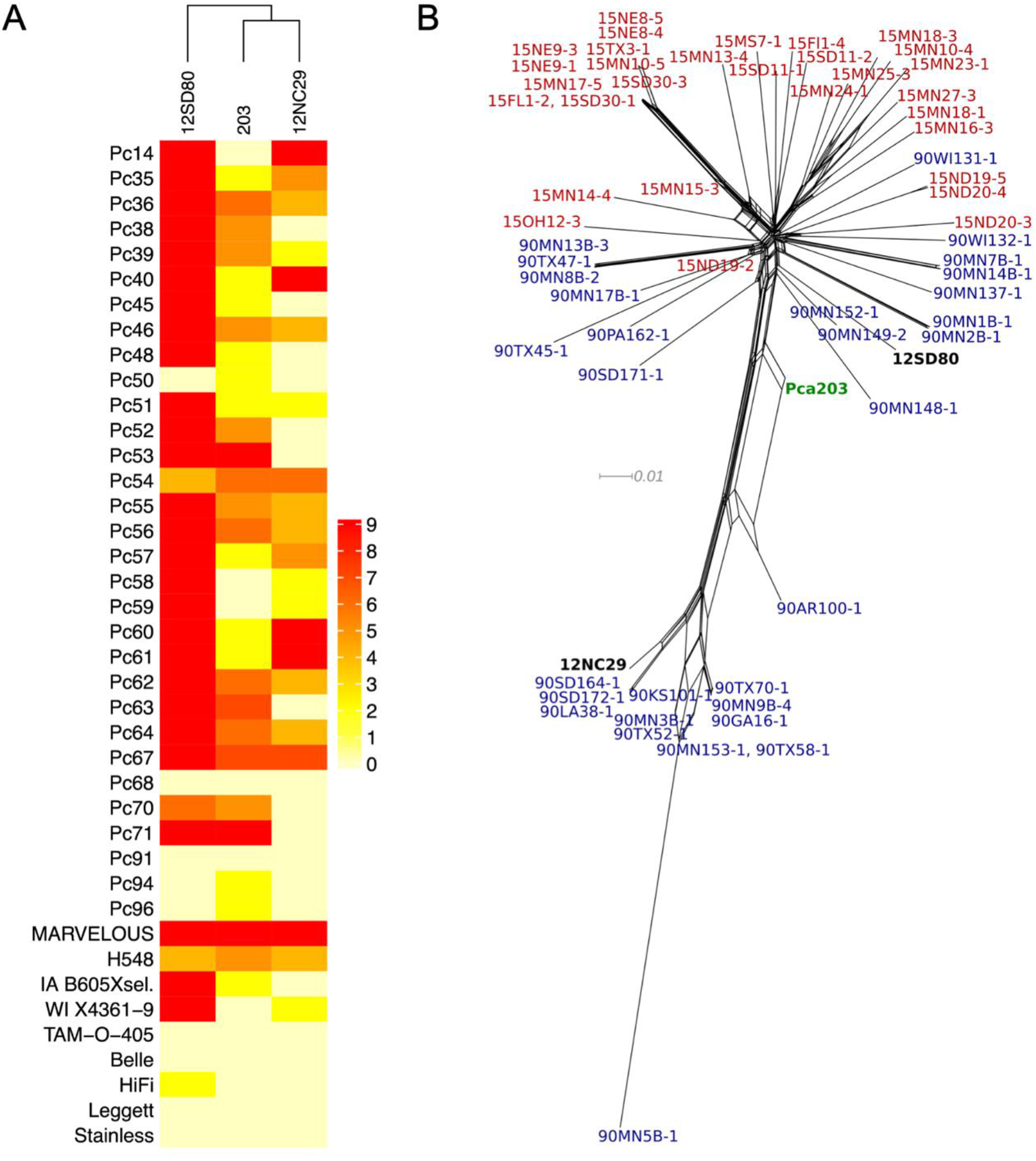
Heatmap of linearized rust scores for *Puccinia coronata* f. sp*. avenae* isolates *Pca*203, 12NC29, and 12SD80 on the North American differential set. Infection scores were converted to a numeric scale (0 = resistance shown in yellow to 9 = susceptibility shown in red) for heatmap generation. Dendrogram (x axis) show hierarchical clustering of isolates with similar virulence patterns. Oat differential lines are shown in the y axis **B)** Neighbor-net network (SplitsTree) of *Pca*203, 12SD80, 12NC29, and additional 60 isolates published previously by Miller et al. (2020). *Pca* isolates derived from 2015 are shown in red whereas isolates collected in 1990 are shown in blue. *Pca*203 is shown in green, and 12NC29 and 12SD80 are displayed in black.

To confirm isolate purity Illumina DNA reads (**Table S1**) from *Pca*203 were mapped to the 12SD80 genome reference (Miller *et al*. 2018), and the allele balance of bi-allelic SNPs was analyzed according to standard methods (Yoshida *et al*. 2013, Li et al. 2019). Results showed the expected binary distribution typical of the presence of two genomes and thus ruled out any potential contaminants from additional *Pca* genotypes (**Figure S2**). These data were combined with SNP data from previously sequenced isolates collected in 1990 and 2015 and a neighbor-net network analysis (SplitsTree) derived from 974,924 variant sites to understand the genetic relationship of *Pca*203 to these temporally distant *Pca* populations. This analysis placed *Pca*203 in a central position between the two major clades detected among populations (**Figure 1B**). This finding is consistent with *Pca*203 being part of the ancestral North American *Pca* population from which these modern populations are derived, although this is difficult to determine due to the lack of historical population samples spanning years prior 1990.

### Genome assembly, curation and construction of chromosomes

In total, 51 Gb of PacBio data (~100x coverage of the *Pca* genome) (**Table S1**) were assembled and polished with 19 Gb of Illumina short reads (~58x coverage of the *Pca* genome). After removal of contaminant and low coverage contigs this resulted in an assembly of 658 contigs adding to a total size of 206.4 Mb (**Table S2**). Given that the haploid genome assembly size of *Pca* isolates 12SD80 and 12NC29 were estimated at 99.2 and 105.3 Mb, respectively (Miller et al. 2018), these results suggest that the two haplotypes were captured by this approach. Collapsed regions were determined by inspecting the average PacBio read coverage in 1000 bp bins (**Figure S3**). The first large peak at ~177x coverage represents the coverage for most of the haploid or non-collapsed regions. A second small peak near 234x coverage, or double the haploid coverage, likely represents collapsed regions. A total of 1.8 Mb of sequence showed coverage >175x (>1.5 times haploid coverage). Thus, the *Pca*203 assembly only has low numbers of collapsed regions.

The initial *Pca*203 assembly was phased using the NuclearPhaser pipeline (Duan *et al*., 2021), which uses a Hi-C graph approach to phase dikaryon assemblies and includes a step for identifying phase swaps (**Figure 2).** As a first step NuclearPhaser was used to preliminarily assign contigs to the two haplotypes and identify phase swaps using Hi-C contact information. We identified phase swaps in 30 contigs, which were manually corrected by breaking these contigs at the phase switch sites. NuclearPhaser was then used a second time and an additional seven breakpoints were identified in three contigs. After phase correction, the two haplotypes were scaffolded separately into chromosomes using the Hi-C data. Scaffolding resulted in 18 chromosomes in each haplotype, covering 101.696 and 98.492 Mb, with 173 small contigs covering 7.973 Mb remaining unassigned to either haplotype (**Table 1**). Hi-C contact maps were then inspected to identify centromeres in each chromosome of both haplotypes. All centromeres were visible on the Hi-C contact maps for each of the 2*18 chromosome (**Figure 3**), which were ordered according to the numbering of the homologous chromosomes in *Puccinia graminis* f. sp. *tritici* (Li *et al*. 2019). Correct haplotype phasing was confirmed by evaluating the distribution of Hi-C contacts to haplotype A (**Figure 4**). Telomere sequence analysis identified 48 total telomeres of the expected 72, with 20 of 36 in haplotype A and 28 of 36 in haplotype B. The high number of identified telomeres suggest that the entire sequence was acquired for most chromosomes.

**Figure 2.**
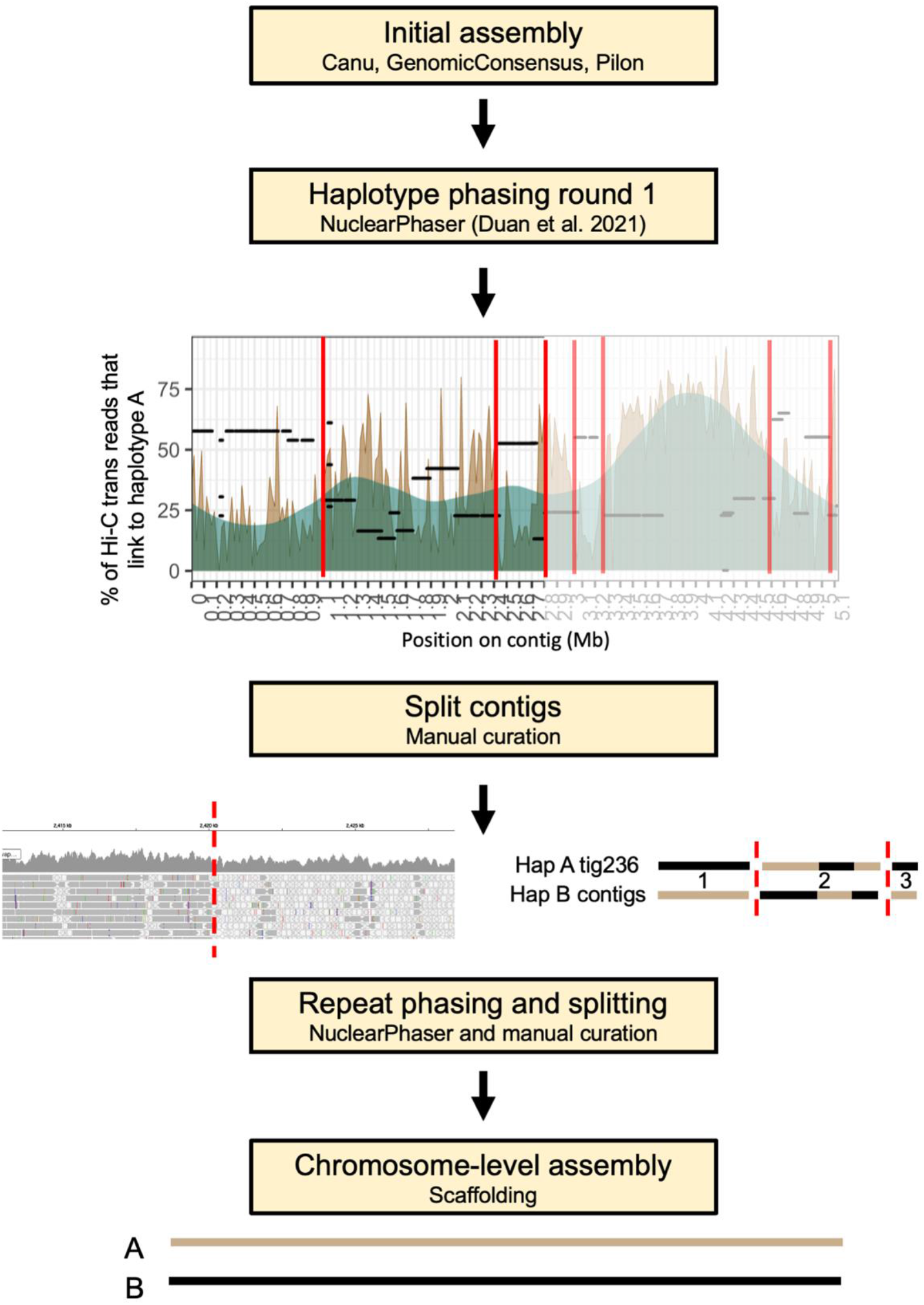
Flowchart illustrating the key steps in the haplotype phasing and chromosome level assembly of *Pca*203. The NuclearPhaser pipeline (Duan *et al*., 2021) was used to create a fully-phased, chromosome-level assembly. First, NuclearPhaser constructs a highly confident subset of the two haplotypes that are expected to reside in separate nuclei and identifies potential phase swaps in the two preliminary haplotype sets. A high proportion of *trans* Hi-C reads should map within the A haplotype, so positions where the proportion drops are flagged as suspect for phase swaps. We inspected these potential phase switch breakpoints using Illumina read mappings. As shown in the figure, a change to high coverage multimapping reads indicates high similarity between regions across haplotypes that may have resulted in a phase swap. After correcting phase swaps, the NuclearPhaser pipeline was used again with the updated genome. Lastly, the two haplotypes were scaffolded separately with Hi-C data into chromosomes.

**Figure 3.**
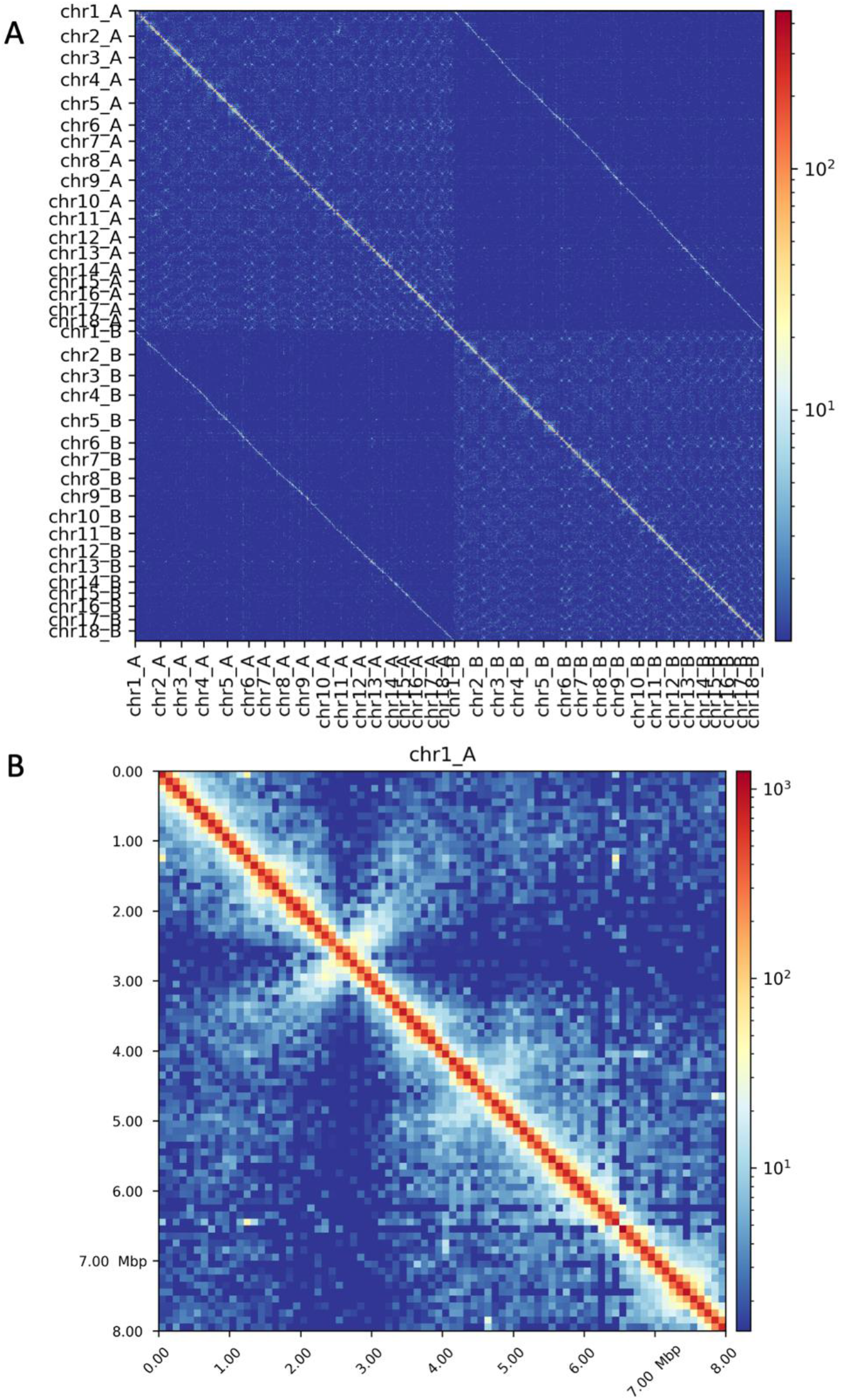
Hi-C contact maps in A) for the *Puccinia coronata* f. sp. *avenae* isolate *Pca*203 A and B haplotypes, and B) for *Pca*203 chromosome 1A in 100 Kb resolution. The two haplotypes exhibit a clear phasing signal, with spurious Hi-C contacts between the haplotypes visible as a weak additional diagonal line in the upper right and lower left corners. The 2*18 centromeres are visible as bowtie-shapes in the contact maps, with chromosome 1A having its centromere located at ~2.5 Mb. Color scale corresponds to the number of contacts in each 100 Kb bin.

**Figure 4.**
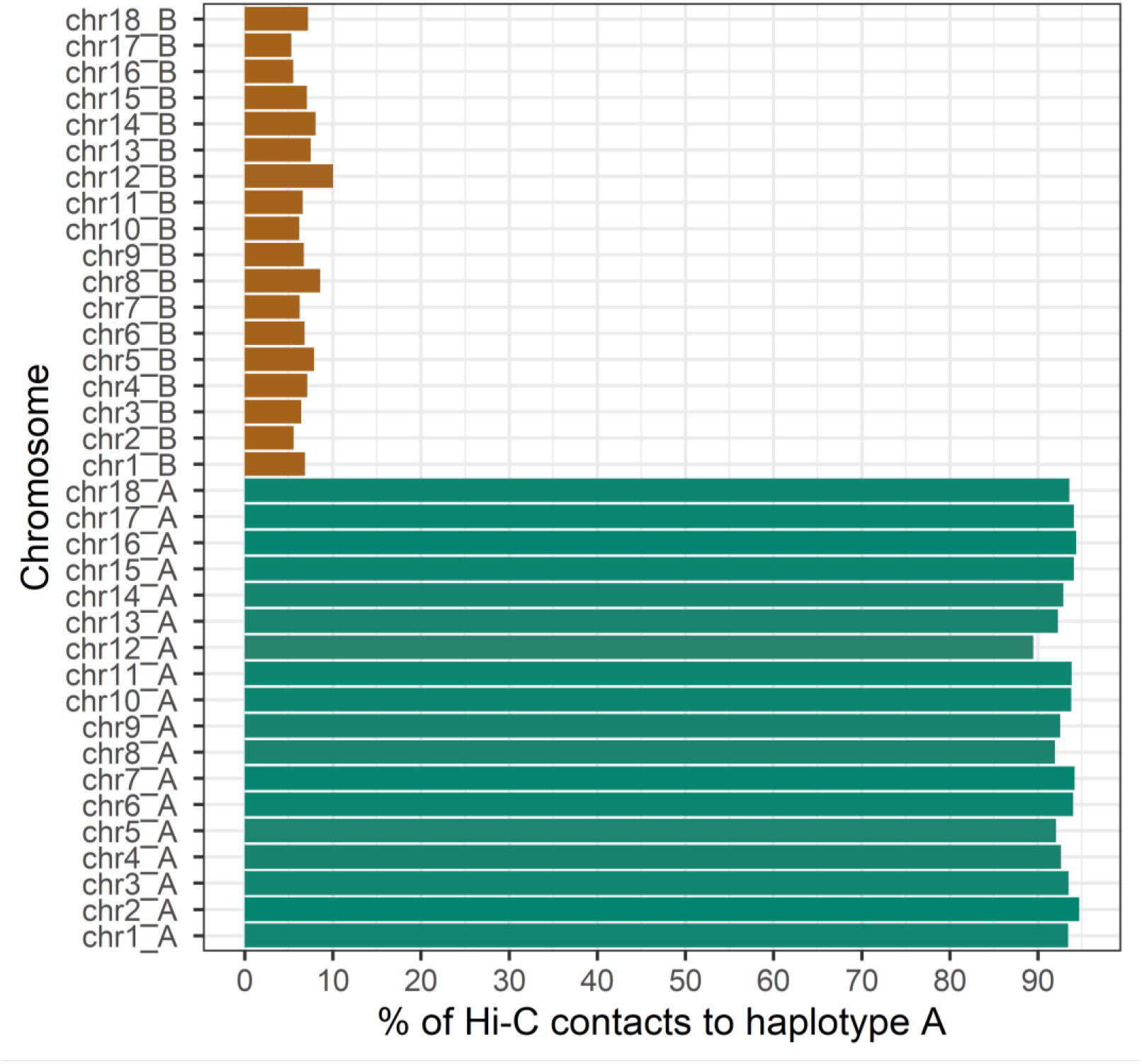
Hi-C *cis* and *trans* contact distribution relative to haplotype A. The haplotype A chromosomes have on average over 90% of their Hi-C links to haplotype A, whereas the haplotype B chromosomes have less than 10% of their Hi-C links to haplotype A.

**Table 1.** Genome assembly and annotation statistics for the chromosome-level nuclear phased genome reference of *Puccinia coronata* f. sp. *avenae* isolate *Pca*203.

|  | 203 Haplotype A | 203 Haplotype B | Unassigned* |
| --- | --- | --- | --- |
| Total size (Mbp) | 101.696 | 98.492 | 7.973 |
| Number of chromosomes | 18 | 18 | 173 contigs |
| Number of telomeres | 20 | 28 | 7 |
| BUSCOs for the combined assembly |  |  |  |
| Complete |  | 93.4% |  |
| Single-copy |  | 9.7% |  |
| Duplicated |  | 83.7% |  |
| Fragmented |  | 2.9% |  |
| Missing |  | 3.7% |  |
| Annotated genes | 17,877 | 17,294 | 1,322 |
| Gene content of genome (%) | 25.0 | 25.2 | 20.3 |
| Number of repeats identified | 146,879 | 141,763 | 10,448 |
| Repeat content of genome (%) | 56.3 | 56.2 | 60.9 |
| Secreted proteins | 1,985 | 1,943 | 108 |
| Predicted effectors | 1,005 | 972 | 52 |

\*Unassigned contigs were not placed in either haplotype.

The completion of the *Pca*203 genome assembly provides the opportunity for comparison to the preexisting high quality references for wheat stem rust (*Pgt*21-0) and wheat leaf rust (*Pt*76). Like the chromosome-level nuclear phased assemblies for *Pgt*21-0 and *Pt*76, the *Pca*203 assembly has 18 chromosome pairs (**Figure 4**). Sequence identity between *Pca*203 and both *Pgt* 21-0 and *Pt*76 is relatively low, as illustrated by dotplot alignments (**Figure 5**). Genome annotation using RNAseq data from *Pca*203, 12SD80, and 12NC29 yielded a total of 36,493 genes, with 17,877 on haplotype A, 17,294 on haplotype B, and the remaining 1,322 on unassigned scaffolds (**Table 1**). Gene space occupies 25.0% on the A haplotype, 25.2% on the B haplotype, and 20.3% of unassigned scaffolds (**Table 1**). In comparison, repeat sequences represent 56.5% of the entire genome, with slightly greater prevalence on the unassigned scaffolds compared to the chromosomes (**Table 1**). Of the 36,524 annotated genes, 4,036 were identified as secreted proteins, and 2,029 were predicted as effectors by EffectorP 3.0 (**Table 1**).

**Figure 5.**
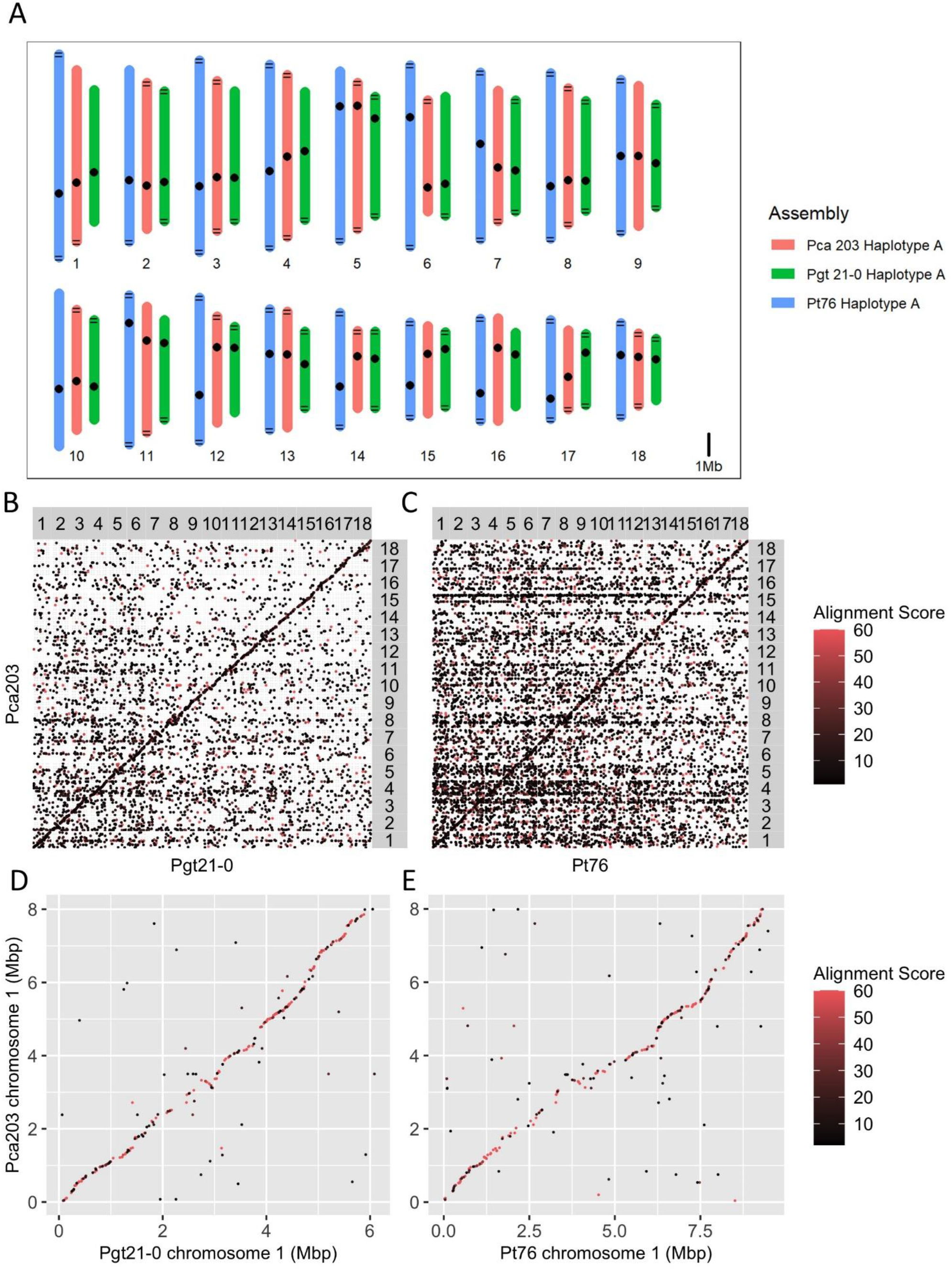
A) Chromosome size comparison between *Puccinia coronata* f. sp. *avenae* isolate *Pca*203 (red), *Puccinia graminis* f. sp. *tritici* isolate *Pgt*21-0 (green), and *Puccinia triticina* isolate *Pt*76 (blue), with dots representing centromeres and black horizontal lines representing identified telomeres. B-E) Dotplot alignment between: B) the isolate *Pca*203 A genome and isolate Pgt 21-0 A genome C) isolate *Pca*203 A genome and isolate *Pt*76 A genome D) isolate *Pca*203 chromosome 1A and isolate *Pgt*21-0 chromosome 1A E) isolate *Pca*203 chromosome 1A and isolate *Pt*76 chromosome 1A. Alignment scores are the mapQ values produced by minimap2; points with MAPQ = 0 are excluded to remove multimapping.

Interestingly, gene synteny among all three cereal rust species is relatively high, as illustrated by the positions of the BUSCO genes shown in **Figure 6**. Compared to *Pgt* 21-0 and *Pt*76, *Pca*203 has slightly more annotated genes than *Pt*76 (31,930) and slightly fewer than *Pgt* 21-0 (38,007). The proportion of the *Pca*203 genome covered by genes is ~ 25% in both the A and B haplotypes, with unplaced contigs being slightly less gene-rich (20%). The gene space in *Pca*203 is comparable to *Pt*76 (24.6%) and lower than that of *Pgt*21-0 (34.9%).

**Figure 6.**
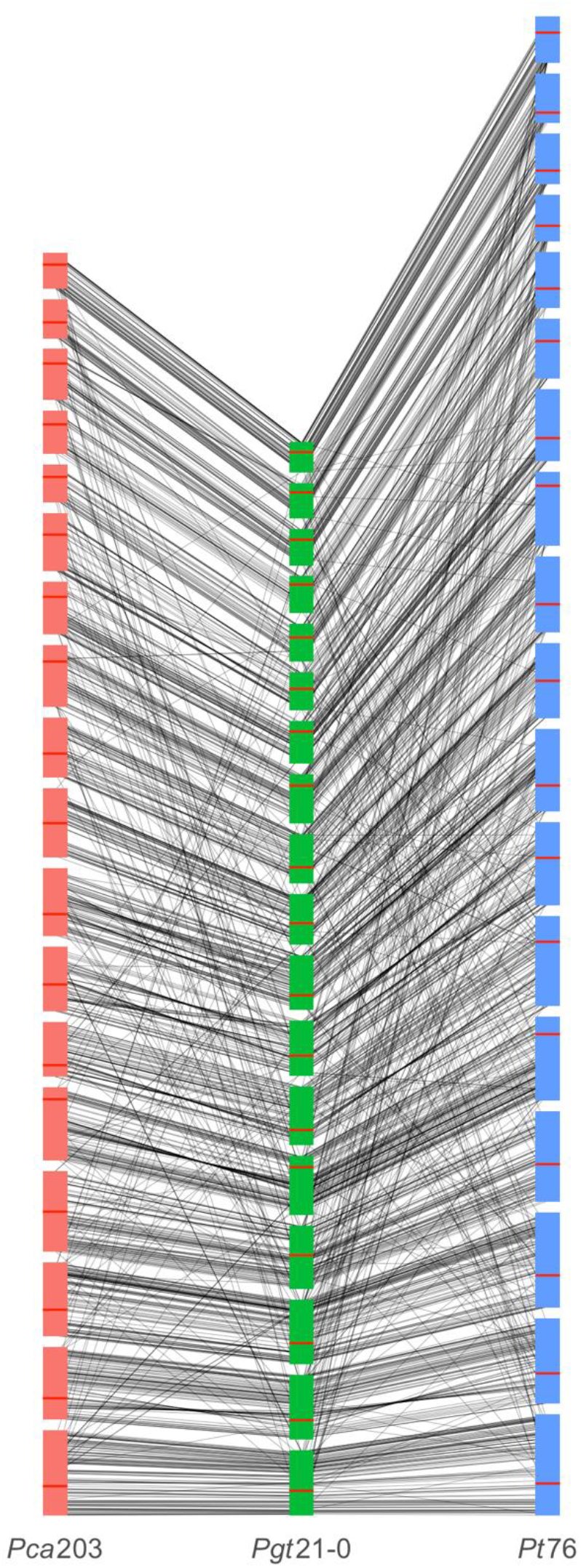
Synteny plot demonstrating connections between BUSCO genes of the *Puccinia coronata* f. sp. *avenae* isolate *Pca*203 haplotype A chromosomes (shown in red) and the haplotype A chromosomes of the cereal rust species *P. graminis* f. sp. *tritici* (*Pgt*21-0, shown in green) and *P. triticina* (*Pt76* shown in blue). Horizontal red lines in chromosomes indicate position of the centromere. Chromosomes are ordered from one to 18 ascending from bottom.

## Conclusions

So far, only a few genome references for rust fungi have been assembled to fully represent phased haplotypes and chromosome sequences. These include the *Pgt*21-0 isolate of the stem rust fungus *P. graminis* f. sp. *tritici* (Li et al., 2019) and the *Pt*76 isolate of the leaf rust fungus *P. triticina* (Duan et al. 2021). Our study introduces the third rust species for which a chromosome-level nuclear phased assembly is available, namely the oat crown rust pathogen, *P. coronata* f. sp. *avenae* (*Pca*). These three *Puccinia* species have 2*18 chromosomes that display high gene synteny despite low overall sequence identity, thus reflecting the close evolutionary history of the species (Aime *et al*. 2018; Aime and McTaggart, 2021).

The chromosome level and nuclear phased assembly of *Pca*203 will enable future pathogenicity studies of this important oat pathogen. At present, no effector gene has been identified for *Pca* and no *R* genes have been isolated from oat or its wild relatives, which is a major bottleneck for disease management strategies. We foresee that access to a gold standard genome reference for *Pca* will accelerate progress in understanding the molecular and genetic basis of the oat crown rust pathosystem. From a plant pathology perspective, the race 203 of *Pca* offers the opportunity to unravel a puzzling question in the field, namely the relationship between genetic resistance to biotrophic pathogens and susceptibility to necrotrophic pathogens. The deployment and widespread use of Victoria and Victoria related oats in the USA as response to oat crown rust epidemics likely caused by *Pca* predominant races such as 203 resulted in significant Victoria Blight epidemics between 1946 and 1948 (Lorang et al., 2007). The fungus *C. victoriae*, causal agent of Victoria Blight, produces a toxin called victorin which is crucial for pathogenicity of *C. victoriae*. Both toxin sensitivity and Victoria blight disease susceptibility are conferred by the gene named *Vb* (Wolpert et al., 2002). The *Vb* gene appears to be genetically linked to the resistance to race 203, conferred by the *Pc2* gene, and it has been suggested that both traits are controlled by the same gene (Wolpert et al., 2002; Welsh *et al*. 1954; Mayama *et al*. 1995). The effector that would be recognized by *Pc2*, *AvrPc2*, has not been identified yet; however, the *Pca*203 genome assembly may aid in the identification as the *AvrPc2* sequence should be represented in the assembly. Given the extensive characterization of Victorin and additional genetic components (Wolpert *et al*. 1985, 1989; Lorang et al., 2007; Kessler *et al*. 2020) dictating the outcome of this plant pathogen interaction, the identification of *AvrPc2* is an important piece to understand the trade-offs of genetic resistance. As highlighted by the virulence profile of *Pca*203, other effectors recognized by dozens of immunoreceptors in oat should also be present in this assembly.

In the past few decades, *Pca* has drastically shifted towards a wider spectrum of virulence (Miller et al. 2020). While this situation was analyzed in depth within US-derived populations, researchers from all over the world have made similar observations. Thus, the development of virulence markers and robust surveillance activities with capacity for large numbers of samples seems particularly critical for pathosystems like oat crown rust which display rapid pathogen evolution and need for durable genetic resistance. This genome reference will aid in the discovery of effector allelic variants, which will enable researchers to develop such tools for virulence monitoring strategies.

## Acknowledgements

We thank Jakob Riddle and Roger Caspers at USDA-ARS CDL for technical assistance. This work utilized computational resources at CSIRO and the Minnesota Supercomputing Institute (https://www.msi.umn.edu).

## Author contributions

M.F., J.S., P.N.D., S.F.K. and B.J.S. conceived and directed the study. E.C.H., F.L., E.S.N., M.F. and S.D. collected samples, isolated DNA/RNA and prepared biological materials for sequencing and other service providers. E.C.H, T.H, E.G. and J.S conducted data analysis. All authors contributed to data interpretation. E.C.H, P.N.D, J.S and M.F. wrote the manuscript. All authors edited the manuscript.

---

**Table S1.** Sequencing statistics for raw reads used in the *Puccinia coronata* f. sp. *avenae* isolate *Pca*203 genome assembly and annotation.

| SRA | Type | Sequencer | Total Raw Reads | Total length of reads (bp) | Average coverage |
| --- | --- | --- | --- | --- | --- |
|  | Genomic DNA | Pacbio Sequel II | 9,041,864 | 28,267,975,258 | 135.8 |
|  | Genomic DNA | Illumina NovaSeq S2 | 50,934,284 | 7,691,076,884 | 36.9 |
|  | RNA (2dpi) | Illumina NextSeq | 149,868,264 | 11,389,988,064 | - |
|  | RNA (5dpi) | Illumina NextSeq | 161,754,544 | 12,293,345,344 | - |
|  | Hi-C Library | Illumina | 130,793,620 | 19,619,043,000 | 94.2 |

**Table S2.**
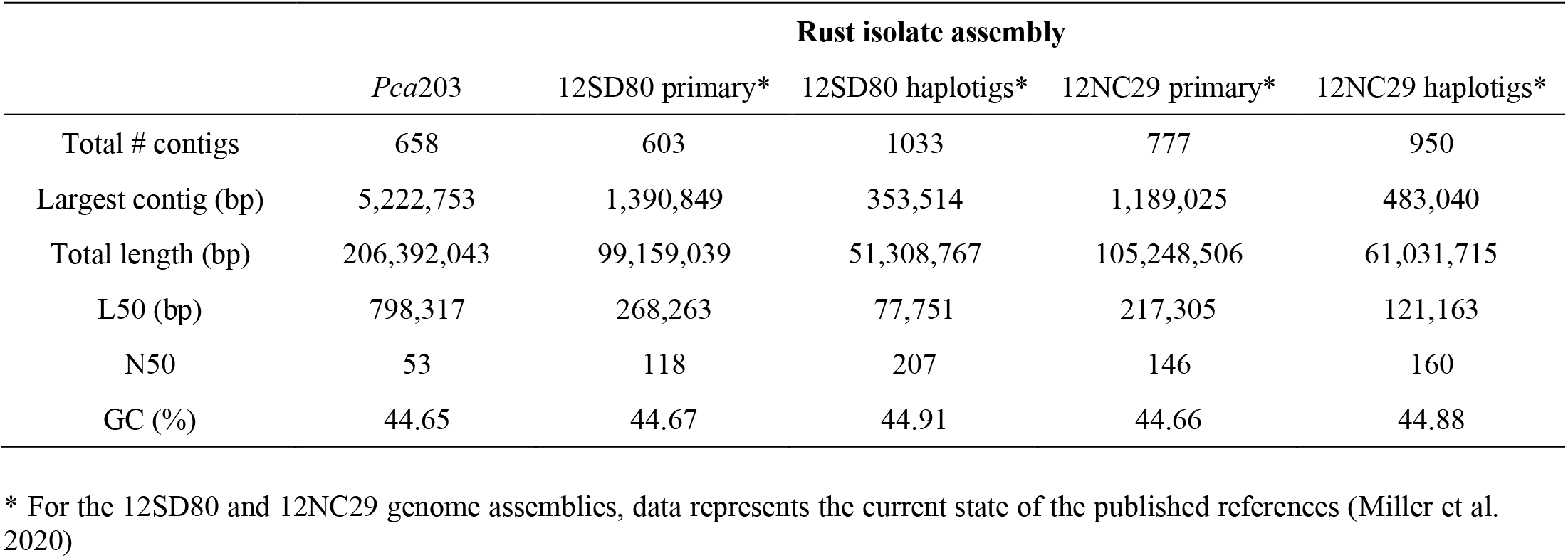
Genome assembly statistics for *Puccinia coronata* f. sp. avenae isolate *Pca*203 after polishing and removal of contaminants and mitochondrial sequences and before scaffolding and haplotype phasing. Genome assembly statistics of 12SD80 and 12NC29 were included for comparison purposes.

**Figure S1.**
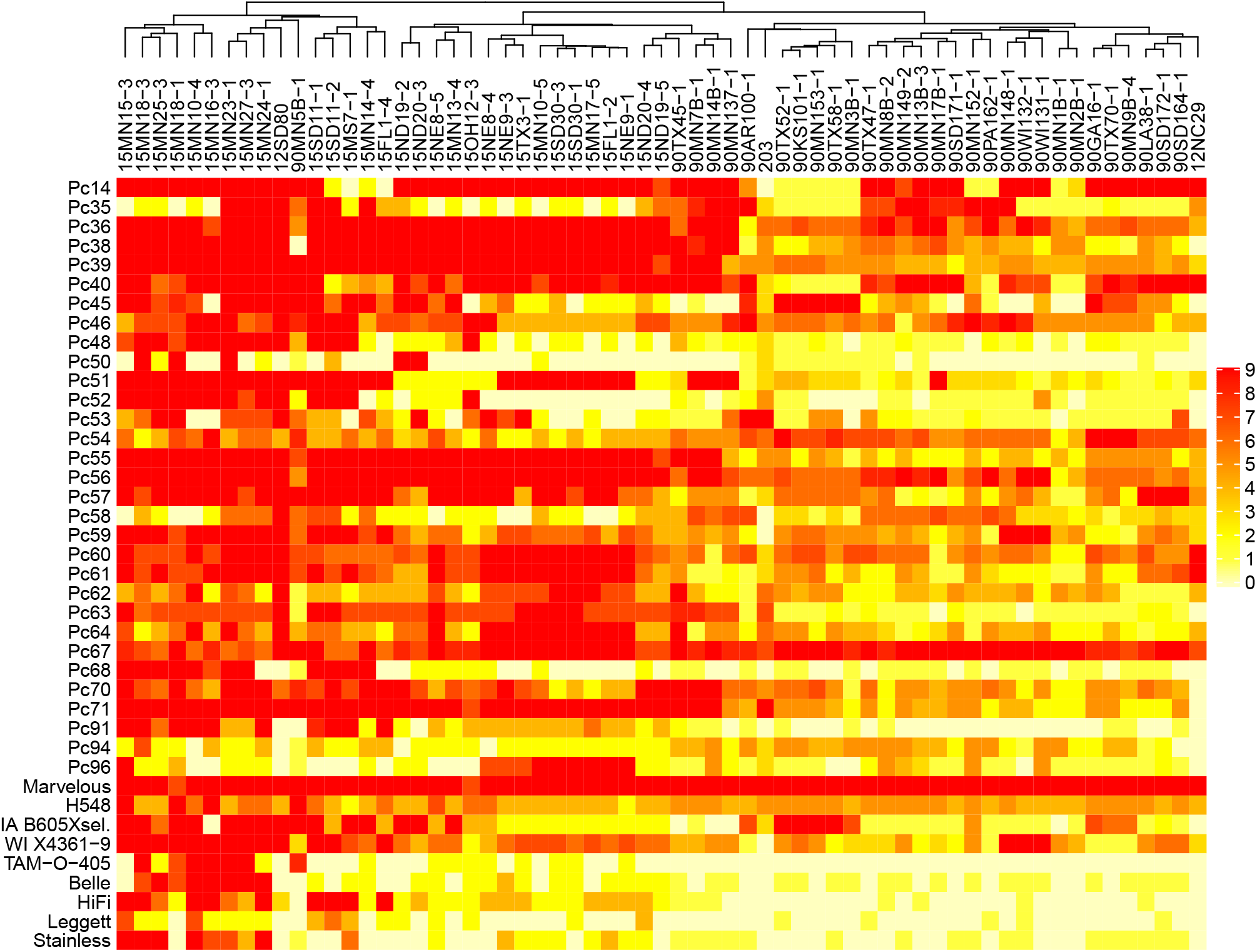
Heatmap of linearized rust scores on the North American differential set for *Puccinia coronata* f. sp. *avenae* isolates *Pca*203, 12SD80, 12NC29, 30 isolates from 1990 and 30 isolates from 2015 as previously published in Miller et al. (2020). Infection scores were converted to a numeric scale (0 = resistance shown in yellow to 9 = susceptibility shown in red) for heatmap generation. Dendrogram (x-axis) shows hierarchical clustering of isolates with similar virulence patterns. Oat differential lines are shown in the y-axis.

**Figure S2.**
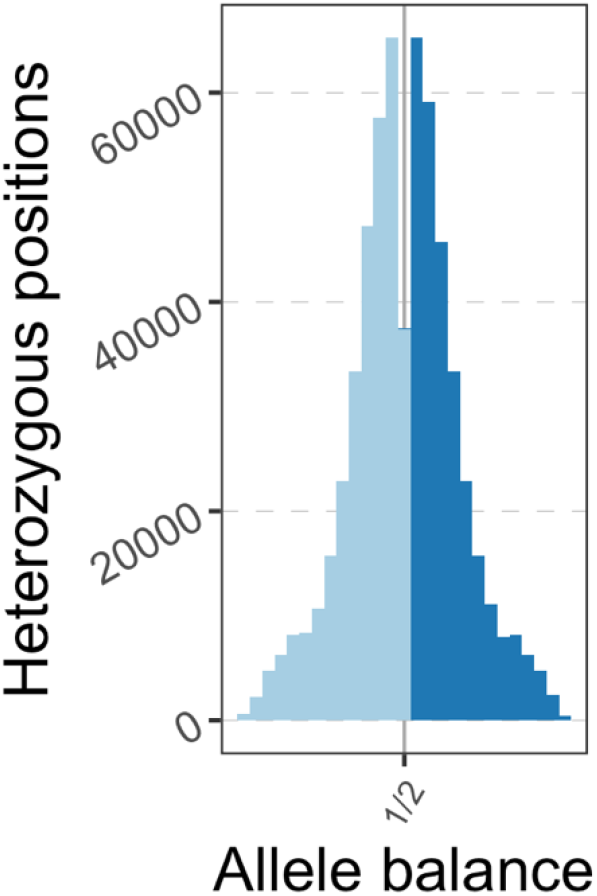
Allele balance plot of variants in *Puccinia coronata* f. sp. *avenae* isolate *Pca*203 when mapped against the 12SD80 reference genome.

**Figure S3.**
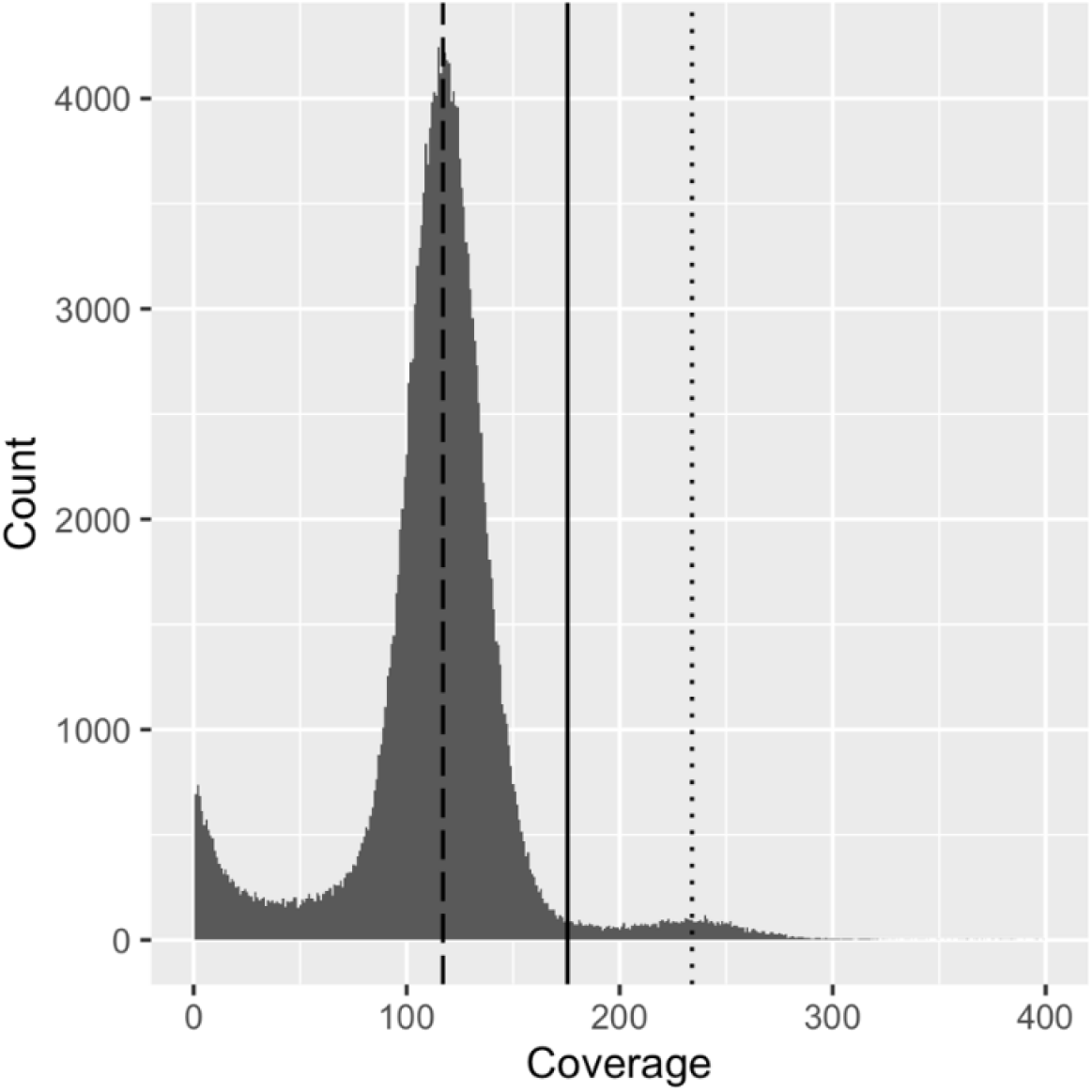
Histogram of the average coverage in 1000 bp bins across the cleaned *Puccinia coronata* f. sp. *avenae* isolate *Pca*203 genome assembly. Dashed line (x = 117) represents the coverage mode, which represents the haploid coverage for the assembly. The dotted line represents double the haploid coverage (x = 234). The solid line (x = 175) is the cutoff used for determining whether a region was considered collapsed.

## Literature Cited

Aime MC, Bell CD, Wilson AW. 2018. Deconstructing the evolutionary complexity between rust fungi (Pucciniales) and their plant hosts. Studies in Mycology. 89:143–152. doi:10.1016/j.simyco.2018.02.002.

Aime MC, McTaggart AR. 2021. A higher-rank classification for rust fungi, with notes on genera. Fungal Systems and Evolution. 7:21–47. doi:10.3114/fuse.2021.07.02.

Bolger AM, Lohse M, Usadel B. 2014. Trimmomatic: A flexible trimmer for Illumina Sequence Data. Bioinformatics.

Cabanettes F, Klopp C. 2018. D-GENIES: dot plot large genomes in an interactive, efficient and simple way. PeerJ. 6:e4958. doi:10.7717/peerj.4958.

Camacho C, Coulouris G, Avagyan V, Ma N, Papadopoulos J, Bealer K, Madden TL. 2009. BLAST+: architecture and applications. BMC Bioinformatics. 10(1):421. doi:10.1186/1471-2105-10-421.

Cantu D, Govindarajulu M, Kozik A, Wang M, Chen X, Kojima KK, Jurka J, Michelmore RW, Dubcovsky J. 2011. Next Generation Sequencing Provides Rapid Access to the Genome of *Puccinia striiformis* f. sp. *tritici*, the Causal Agent of Wheat Stripe Rust. PLOS ONE. 6(8):e24230. doi:10.1371/journal.pone.0024230.

Chang TD, Sadanaga K. 1964. Crosses of Six Monosomics in *Avena sativa* L. With Varieties, Species, and Chlorophyll Mutants1. Crop Science. 4(6):cropsci1964.0011183X000400060012x. doi:10.2135/cropsci1964.0011183X000400060012x.

Chen S, Zhou Y, Chen Y, Gu J. 2018. fastp: an ultra-fast all-in-one FASTQ preprocessor. Bioinformatics. 34(17):i884–i890. doi:10.1093/bioinformatics/bty560.

Chong J, Leonard KJ, Salmeron JJ. 2000. A North American System of Nomenclature for *Puccinia coronata* f. sp. *avenae*. Plant Disease. 84(5):580–585. doi:10.1094/PDIS.2000.84.5.580.

Cuomo CA, Bakkeren G, Khalil HB, Panwar V, Joly D, Linning R, Sakthikumar S, Song X, Adiconis X, Fan L, et al. 2017. Comparative Analysis Highlights Variable Genome Content of Wheat Rusts and Divergence of the Mating Loci. G3 Genes|Genomes|Genetics. 7(2):361–376. doi:10.1534/g3.116.032797.

Danecek P, Auton A, Abecasis G, Alber CA, Banks E, DePristo MA, Handsaker R, Lunter G, Marth G, Sherry ST, et al. 2011. The Variant Call Format and VCFtools. Bioinformatics.

Dodds PN, Rathjen JP. 2010. Plant immunity: towards an integrated view of plant-pathogen interactions. Nature Reviews Genetics. 11(8):539–548. doi:10.1038/nrg2812.

Duan H, Jones AW, Hewitt T, Mackenzie A, Hu Y, Sharp A, Lewis D, Mago R, Upadhyaya NM, Rathjen JP, et al. 2021. Identification and correction of phase switches with Hi-C data in the Nanopore and HiFi chromosome-scale assemblies of the dikaryotic leaf rust fungus Puccinia triticina. https://www.biorxiv.org/content/10.1101/2021.04.28.441890v1.

Figueroa M, Dodds PN, Henningsen EC. 2020. Evolution of virulence in rust fungi — multiple solutions to one problem. Current Opinion in Plant Biology. 56:20–27. doi:10.1016/j.pbi.2020.02.007.

Figueroa M, Upadhyaya NM, Sperschneider J, Park RF, Szabo LJ, Steffenson B, Ellis JG, Dodds PN. 2016. Changing the Game: Using Integrative Genomics to Probe Virulence Mechanisms of the Stem Rust Pathogen *Puccinia graminis* f. sp. tritici. Frontiers in Plant Science. 7:205. doi:10.3389/fpls.2016.00205.

Fleischmann G. 1967. Virulence of uredial and aecial isolates of *Puccinia coronata* f. sp. avenae identified in Canada from 1952 to 1966. Canadian Journal of Botany. 45:1693–1701.

Fleischmann G, Baker FJ. 1971. Oat crown rust differentiation: replacement of the standard differential varieties with a new set of single resistance gene lines derived from *A. sterilis*. Canadian Journal of Plant Pathology. 25:97–108.

Flynn JM, Hubley R, Goubert C, Rosen J, Clark AG, Feschotte C, Smit AF. 2020. RepeatModeler2 for automated genomic discovery of transposable element families. Proc Natl Acad Sci USA. 117(17):9451. doi:10.1073/pnas.1921046117.

Garrison E, Marth G. 2012. Haplotype-based variant detection from short-read sequencing. arXiv preprint. (arXiv:1207.3907).

Ghurye J, Pop M, Koren S, Bickhart D, Chin C-S. 2017. Scaffolding of long read assemblies using long range contact information. BMC Genomics. 18(1):527. doi:10.1186/s12864-017-3879-z.

Ghurye J, Rhie A, Walenz BP, Schmitt A, Selvaraj S, Pop M, Phillippy AM, Koren S. 2019. Integrating Hi-C links with assembly graphs for chromosome-scale assembly. PLOS Computational Biology. 15(8):e1007273. doi:10.1371/journal.pcbi.1007273.

Grabherr MG, Haas BJ, Yassour M, Levin JZ, Thompson DA, Amit I, Adiconis X, Fan L, Raychowdhury R, Zeng Q, et al. 2011. Full-length transcriptome assembly from RNA-Seq data without a reference genome. Nat Biotechnol. 29(7):644–652. doi:10.1038/nbt.1883.

Gu Z, Eils R, Schlesner M. 2016. Complex heatmaps reveal patterns and correlations in multidimensional genomic data. Bioinformatics. 32(18):2847–2849. doi:10.1093/bioinformatics/btw313.

Gu Z, Gu L, Eils R, Schlesner M, Brors B. 2014. circlize implements and enhances circular visualization in R. Bioinformatics. 30(19):2811–2812. doi:10.1093/bioinformatics/btu393.

Gurevich A, Saveliev V, Vyahhi N, Tesler G. 2013. QUAST: quality assessment tool for genome assemblies. Bioinformatics. 29(8):1072–1075. doi:10.1093/bioinformatics/btt086.

Huson DH, Bryant D. 2006. Application of Phylogenetic Networks in Evolutionary Studies. Molecular Biology and Evolution. 23(2):254–267. doi:10.1093/molbev/msj030.

Kessler SC, zhang X, McDonald MC, Gilchrist CLM, Lin Z, Rightmyer A, Solomon PS, Turgeon BG, Chooi Y-H. 2020. Victorin, the host-selective cyclic peptide toxin from the oat pathogen *Cochliobolus victoriae*, is ribosomally encoded. PNAS. 117(39):24243–24250. doi:10.1073/pnas.2010573117.

Kim D, Paggi JM, Park C, Bennett C, Salzberg SL. 2019. Graph-based genome alignment and genotyping with HISAT2 and HISAT-genotype. Nature Biotechnology. 37(8):907–915. doi:10.1038/s41587-019-0201-4.

Koren S, Walenz BP, Berlin K, Miller JR, Bergman NH, Phillippy AM. 2017. Canu: scalable and accurate long-read assembly via adaptive k-mer weighting and repeat separation. Genome Research. 27:722–736. doi:10.1101/gr.215087.116.

Krogh A, Larsson B, von Heijne G, Sonnhammer ELL. 2001. Predicting transmembrane protein topology with a hidden markov model: application to complete genomes. Journal of Molecular Biology. 305(3):567–580. doi:10.1006/jmbi.2000.4315.

Li F, Upadhyaya NM, Sperschneider J, Matny O, Nguyen-Phuc H, Mago R, Raley C, Miller ME, Silverstein KAT, Henningsen E, et al. 2019. Emergence of the Ug99 lineage of the wheat stem rust pathogen through somatic hybridisation. Nature Communications. 10(1):5068. doi:10.1038/s41467-019-12927-7.

Li H. 2018. Minimap2: pairwise alignment for nucleotide sequences. Bioinformatics. 34(18):3094–3100. doi:10.1093/bioinformatics/bty191.

Li H, Durbin R. 2009. Fast and accurate short read alignment with Burrows-Wheeler Transform. Bioinformatics. 25:1754–1760.

Li H, Handsaker B, Wysoker A, Fennell T, Ruan J, Homer N, Marth G, Abecasis G, Durbin R, 1000 Genome Project Data Processing Subgroup. 2009. The Sequence Alignment/Map format and SAMtools. Bioinformatics. 25(16):2078–2079. doi:10.1093/bioinformatics/btp352.

Lorang JM, Sweat TA, Wolpert TJ. 2007. Plant disease susceptibility conferred by a “resistance” gene. Proc Natl Acad Sci USA. 104(37):14861. doi:10.1073/pnas.0702572104.

Mayama S, Bordin APA, Morikawa T, Tanpo H, Kato H. 1995. Association of avenalumin accumulation with co-segregation of victorin sensitivity and crown rust resistance in oat lines carrying the *Pc*-2 gene. Physiological and Molecular Plant Pathology. 46(4):263–274. doi:https://doi.org/10.1006/pmpp.1995.1021.

Miller ME, Nazareno ES, Rottschaefer SM, Riddle J, Dos Santos Pereira D, Li F, Nguyen-Phuc H, Henningsen EC, Persoons A, Saunders DGO, et al. 2021. Increased virulence of *Puccinia coronata* f. sp. *avenae* populations through allele frequency changes at multiple putative *Avr* loci. PLOS Genetics. 16(12):e1009291. doi:10.1371/journal.pgen.1009291.

Miller ME, Zhang Y, Omidvar V, Sperschneider J, Schwessinger B, Raley C, Palmer JM, Garnica D, Upadhyaya N, Rathjen J, et al. 2018. *De Novo* Assembly and Phasing of Dikaryotic Genomes from Two Isolates of *Puccinia coronata* f. sp. *avenae*, the Causal Agent of Oat Crown Rust. mBio. 9(1):e01650–17. doi:10.1128/mBio.01650-17.

Murphy HC, Meehan F. 1946. Reaction of oat varieties to a new species of *Helminthosporium*. Phytopathology. 36(5):407.

Nazareno ES, Li F, Smith M, Park RF, Kianian SF, Figueroa M. 2018. *Puccinia coronata* f. sp. *avenae:* a threat to global oat production. Molecular Plant Pathology. 19(5):1047–1060. doi:10.1111/mpp.12608.

Omidvar V, Dugyala S, Li F, Rottschaefer SM, Miller ME, Ayliffe M, Moscou MJ, Kianian SF, Figueroa M. 2018. Detection of race-specific resistance againt *Puccinia coronata* f. sp. *avenae* in *Brachypodium* species. Phytopathology. 108(12):1443–1454. doi:10.1094/PHYTO-03-18-0084-R.

Palmer JM, Stajich J. 2020. funannotate v1.7.4.

Petersen TN, Brunak S, von Heijne G, Nielsen H. 2011. SignalP 4.0: discriminating signal peptides from transmembrane regions. Nature Methods. 8(10):785–786. doi:10.1038/nmeth.1701.

Ramírez F, Bhardwaj V, Arrigoni L, Lam KC, Grüning BA, Villaveces J, Habermann B, Akhtar A, Manke T. 2018. High-resolution TADs reveal DNA sequences underlying genome organization in flies. Nature Communications. 9(1):189. doi:10.1038/s41467-017-02525-w.

Servant N, Varoquaux N, Lajoie BR, Viara E, Chen C-J, Vert J-P, Heard E, Dekker J, Barillot E. 2015. HiC-Pro: an optimized and flexible pipeline for Hi-C data processing. Genome Biology. 16(1):259. doi:10.1186/s13059-015-0831-x.

Sperschneider J, Dodds PN. 2021 Jan 1. EffectorP 3.0: prediction of apoplastic and cytoplasmic effectors in fungi and oomycetes. bioRxiv.:2021.07.28.454080. doi:10.1101/2021.07.28.454080.

Stamatakis A. 2014. RAxML version 8: a tool for phylogenetic analysis and post-analysis of large phylogenies. Bioinformatics. 30(9):1312–1313. doi:10.1093/bioinformatics/btu033.

Stoa TE, Swallers CM. 1950. Keeping Up-to-Date on Oats. NDSU Agricultural Experiment Station Bimonthly Bulletin. 12(4):9.

Upadhyaya NM, Mago R, Panwar V, Hewitt T, Luo M, Chen J, Sperschneider J, Nguyen-Phuc H, Wang A, Ortiz D, et al. 2021. Genomics accelerated isolation of a new stem rust avirulence gene–wheat resistance gene pair. Nature Plants. 7(9):1220–1228. doi:10.1038/s41477-021-00971-5.

USDA-ARS CDL. 2014. 2014 Oat Loss to Rust. https://www.ars.usda.gov/ARSUserFiles/50620500/Smallgrainlossesduetorust/2014loss/2014oatloss.pdf

Walker BJ, Abeel T, Shea T, Priest M, Abouelliel A, Sakthikumar S, Cuomo CA, Zeng Q, Wortman J, Young SK, et al. 2014. Pilon: An Integrated Tool for Comprehensive Microbial Variant Detection and Genome Assembly Improvement. PLOS ONE. 9(11):e112963. doi:10.1371/journal.pone.0112963.

Waterhouse RM, Seppey M, Simão FA, Manni M, Ioannidis P, Klioutchnikov G, Kriventseva EV, Zdobnov EM. 2018. BUSCO Applications from Quality Assessments to Gene Prediction and Phylogenomics. Molecular Biology and Evolution. 35(3):543–548. doi:10.1093/molbev/msx319.

Welsh JN, Peturson B, Machacek JE. 1954. Associated inheritance of reaction to races of crown rust, *Puccinia coronata avenae* Erikss., and to Victoria blight, *Helminthosporium victoriae* M. and M., in oats. Can J Bot. 32(1):55–68. doi:10.1139/b54-008.

Winter DJ, Ganley ARD, Young CA, Liachko I, Schardl CL, Dupont P-Y, Berry D, Ram A, Scott B, Cox MP. 2018. Repeat elements organise 3D genome structure and mediate transcription in the filamentous fungus *Epichloë festucae*. PLOS Genetics. 14(10):e1007467. doi:10.1371/journal.pgen.1007467.

Wolff J, Bhardwaj V, Nothjunge S, Richard G, Renschler G, Gilsbach R, Manke T, Backofen R, Ramírez F, Grüning BA. 2018. Galaxy HiCExplorer: a web server for reproducible Hi-C data analysis, quality control and visualization. Nucleic Acids Research. 46(W1):W11–W16. doi:10.1093/nar/gky504.

Wolff J, Rabbani L, Gilsbach R, Richard G, Manke T, Backofen R, Grüning BA. 2020. Galaxy HiCExplorer 3: a web server for reproducible Hi-C, capture Hi-C and single-cell Hi-C data analysis, quality control and visualization. Nucleic Acids Research. 48(W1):W177–W184. doi:10.1093/nar/gkaa220.

Wolpert TJ, Dunkle LD, Ciuffetti LM. 2002. Host-selective toxins and avirulence determinants: What’s in a Name? Annu Rev Phytopathol. 40(1):251–285. doi:10.1146/annurev.phyto.40.011402.114210.

Wolpert TJ, Macko V. 1989. Specific binding of victorin to a 100-kDa protein from oats. Proc Natl Acad Sci U S A. 86(11):4092–4096. doi:10.1073/pnas.86.11.4092.

Wolpert TJ, Macko V, Acklin W, Jaun B, Seibl J, Meili J, Arigoni D. 1985. Structure of victorin C, the major host-selective toxin from *Cochliobolus victoriae*. Experientia. 41(12):1524–1529. doi:10.1007/BF01964789.

Yoshida K, Schuenemann VJ, Cano LM, Pais M, Mishra B, Sharma R, Lanz C, Martin FN, Kamoun S, Krause J, et al. 2013. The rise and fall of the *Phytophthora infestans* lineage that triggered the Irish potato famine. eLife. 2:e00731. doi:10.7554/eLife.00731.

Yu G, Smith DK, Zhu H, Guan Y, Lam TT-Y. 2017. ggtree: an r package for visualization and annotation of phylogenetic trees with their covariates and other associated data. Methods in Ecology and Evolution. 8(1):28–36. doi:10.1111/2041-210X.12628.

